# Cell adhesion factors during adolescence support amygdalo-cortical connections and flexible action later in life

**DOI:** 10.1101/2021.07.12.452062

**Authors:** Henry W. Kietzman, Lauren P. Shapiro, Gracy Trinoskey-Rice, Jidong Guo, Shannon L. Gourley

## Abstract

Adolescent brain development is characterized by dramatic neuronal remodeling in the prefrontal cortex. This plasticity is presumed to act in part to “set the stage” for prefrontal cortical function in adulthood, but causal relationships have largely not been verified. Integrins are cell adhesion factors that provide a link between the extracellular matrix and the intracellular actin cytoskeleton. We find that β1-integrin presence in the prelimbic subregion of the prefrontal cortex (PL) during adolescence, but not adulthood, is necessary for adult mice to select actions based on reward likelihood and value. These behaviors require coordinated limbic-frontal-striatal circuits. We identified projections from the basolateral amygdala (BLA) to PL as being necessary for mice to express learned response strategies. We then visualized adolescent PL neurons receiving input from the BLA and projecting to the dorsomedial striatum (DMS), a primary striatal output by which the PL controls reward-related behavior. These projection-defined neurons had a more “adult-like” morphology relative to a general population of layer V PL neurons. β1-integrin loss caused the overexpression of stubby-type dendritic spines at the expense of more mature spines, a phenotype not observed when β1-integrins were silenced before or after adolescence. Together, these experiments localize β1-integrin-mediated cell adhesion activity within a developing di-synaptic circuit that coordinates flexible action.

## Introduction

Neurodevelopment during adolescence is characterized by dramatic structural plasticity within the prefrontal cortex (PFC), involving both the elimination of dendritic spines and the stabilization of other spines^1, 2^. Early-life development is the subject of intense investigation, but mechanisms underlying adolescent brain development are still unclear – even despite ∼50% of “adult” mental health disorders initially presenting during adolescence^3^. Further, structural irregularities in the PFC are linked to the cognitive symptomatology of nearly every major neurodevelopmental disorder^4-6^. Thus, investigating factors that govern *structural stability* – referring here to the process by which dendritic spines are retained and escape pruning – is a crucial step in understanding neurodevelopment during adolescence and potentially, disease etiologies.

Integrins are heterodimeric transmembrane cell adhesion receptors that provide a link between the extracellular matrix (ECM) and the intracellular actin cytoskeleton. Integrins, composed of a ligand-binding α subunit and a β subunit that activates intracellular signaling cascades, respond to ECMs, either to maintain structural integrity and functionality, or to promote tissue differentiation and development^7^. In early life, β1-subunit-containing integrin receptors promote neurite outgrowth^8^ and axon guidance^9^. Investigations focused on hippocampal CA1 suggest that during adolescence, β1-integrins may fulfill a distinct function: stabilizing dendrites and synapses^10^, thereby supporting long-term potentiation^11^. Levels of the neuronal β1-integrin signaling partners, p190RhoGAP and Rho-kinase 2 (ROCK2), peak in the PFC during adolescence, suggesting that β1-integrin-mediated signaling in adolescence might stabilize PFC neuron structure and function^12^.

Broadly, the PFC, and the prelimbic subregion (PL) in particular, controls the ability of organisms to link actions with their likely outcomes and to flexibly modify behaviors when contingencies change^13, 14^. Here, we report that β1*-*integrin presence in the PL during adolescence, but not later, is necessary for flexible reward-related behavior in adulthood. As such, mice deficient in developmental β1-integrins struggle to: **1)** associate actions with outcomes and **2)** select actions based on outcome value. These behaviors require coordinated limbic-frontal-striatal circuits^14^. We identified projections from the basolateral amygdala (BLA) to PL as being necessary for mice to express learned response strategies. This led us to characterize the dendritic micro-architecture of adolescent layer V PL neurons receiving input from the BLA and projecting to the dorsomedial striatum (DMS), a primary striatal output necessary for goal-oriented action^15^. Surprisingly, these projection-defined neurons had a more “adult-like” morphology relative to a general population of layer V PL neurons. β1-integrin loss caused the overexpression of stubby-type dendritic spines, which are unlikely to contain synapses, at the expense of mature spine types. Together, these experiments indicate that β1-integrin presence during adolescence structurally stabilizes PL neurons positioned within a di-synaptic circuit, providing the structural substrates for flexible action later in life.

## Results

### Cell adhesion during adolescence optimizes flexible action in adulthood

We aimed to understand the function of β1-integrins in the developing postnatal PFC. To reduce β1-integrins, we introduced a CaMKIIα-driven adeno-associated viral vector (AAV) expressing Cre recombinase (Cre) into the PL of transgenic *Itbg1*^*tm1Efu*^ (*Itgb1-*flox) mice. These mice have loxP sites flanking exon 3 of *Itgb1*, the gene that encodes β1-integrins; Cre introduction deletes this exon^16^. Cre was delivered early in life, to reduce β1-integrins starting in adolescence, or for comparison, in adulthood (infusions at P21-24 or ≥P56-60, respectively; ages and viral vector details for all experiments are described in table 1). Fig.1a depicts viral vector spread, comparable across ages; any mice with unilateral or off-target transfection were excluded.

**Figure 1.**
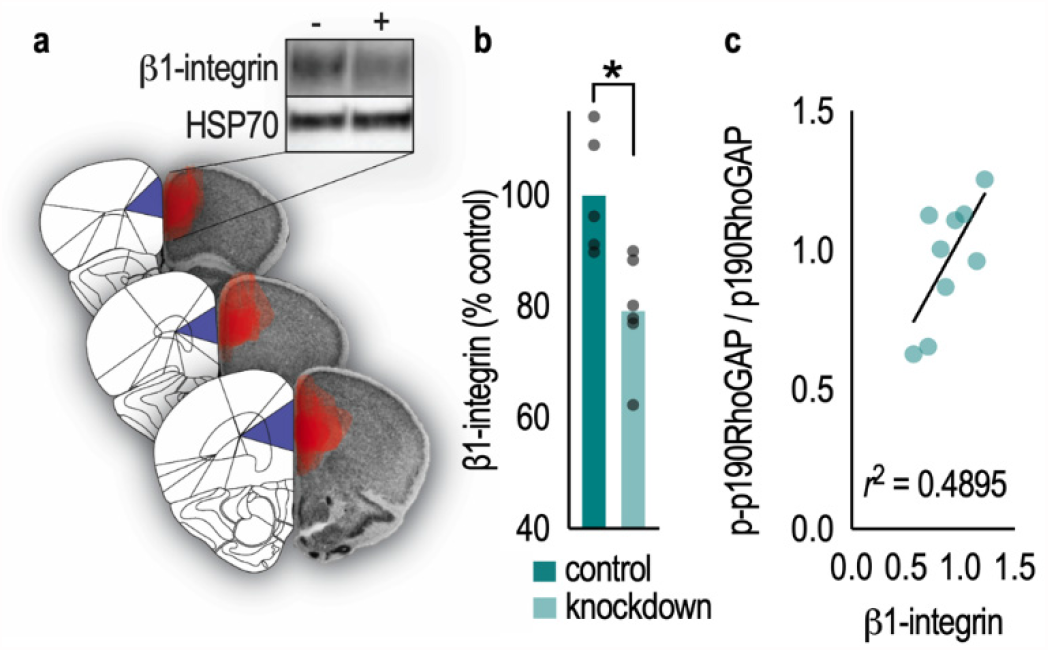
*Itgb1* knockdown reduces β1-integrin protein levels. **a**. CaMKIIα-AAVs ± Cre were infused bilaterally into PL of *Itgb1*-flox mice. (**left**) Images from Allen Brain Atlas (from left: +2.71 mm, +2.22 mm, and +1.98 mm relative to bregma) depicting the PL (purple). (**right**) Overlays of viral vector spread (red) encompassing the PL from mice in this report on images from the Mouse Brain Library^69^. All cohorts exhibited consistent viral vector localization. (**top right**) Representative blots. **b**. Viral-mediated *Itgb1* silencing reduced β1-integrin protein levels in PFC tissue punches by ∼21%. Incomplete protein loss was expected, given that samples contained transduced and unaffected cells, and glial integrins were spared. **c**. β1-integrin protein levels correlated with phosphorylation (activation) of the neuronal substrate p190RhoGAP. Bars represent means, and symbols represent individual mice. \**p*<0.05.

**Table 1.** Genotype, age, and viral vector details for all experiments. Infusions were in the PL unless otherwise indicated in parentheses.

| Fig. | Genotype of mice | Timing of viral vector delivery | Viral vector conditions | Measure |
| --- | --- | --- | --- | --- |
| 1a, 2 | *Itgb1*-flox | P21-24 vs. ≥P56 | *Knockdown:* AAV2/8-CaMKIIα-mCherry+Cre<br>*Control:* AAV2/8-CaMKIIα-mCherry | Behavioral testing |
| 1b-c | *Itgb1*-flox | P21-24 | *Knockdown:* AAV2/8-CaMKIIα-mCherry+Cre<br>*Control:* AAV2/8-CaMKIIα-mCherry | Immunoblotting for $\beta 1$-integrins |
| 3 | wild type | ≥P56 | All mice: rgAAV(pENN)-hSyn-HI-eGFP-Cre-WPRE (PL)<br>AAV5-hSyn-DIO-mCherry+hM4Di (BLA);<br>comparisons were media±CNO or cells±hM4Di | Electrophysiology |
| 4 | wild type | ≥P56 | *Projection-specific inactivation:* rgAAV(pENN)-hSyn-HI-eGFP-Cre-WPRE (PL)<br>AAV5-hSyn-DIO-mCherry+hM4Di (BLA)<br>*Control:* gAAV(pENN)-hSyn-HI-eGFP-Cre-WPRE (PL)<br>AAV5-hSyn-DIO-mCherry (BLA) | Behavioral testing |
| 5, S1 | *Itgb1*-flox/<br>*Thy1*-YFP | P21-24 vs. ≥P56 | *Visualization of projection-defined PL neurons:* rgAAV-hSyn-DIO-mCherry (DMS)<br>AAV1-hSyn-Cre-WPRE-hGH (BLA)<br>*Visualization of general layer V populations:* AAV2/8-CaMKIIα-mCherry | Dendritic spine analysis |
| 6, S1, S2 | *Itgb1*-flox/<br>*Thy1*-YFP | P21-24 vs. ≥P56 | *Adolescent Itgb1 knockdown in projection-defined neurons:*<br>rgAAV-hSyn-DIO-mCherry (DMS)<br>AAV1-hSyn-Cre-WPRE-hGH (BLA)<br>*Adolescent projection-defined control neurons:* rgAAV-hSyn-DIO-mCherry (DMS)<br>AAV1-hSyn-Cre-WPRE-hGH (BLA) (delivered to *Itgb1*-flox<sup>-/-</sup> littermates)<br>*Adolescent- or adult-onset Itgb1 knockdown in general layer V neurons:* AAV2/8-CaMKIIα-mCherry+Cre<br>*Adolescent- or adult-onset general layer V control neurons:* AAV2/8-CaMKIIα-mCherry | Dendritic spine analysis |

Viral-mediated *Itgb1* silencing in young mice reduced β1-integrin protein levels in gross tissue punches by 21% [*t*_(9)_=3.30, *p*=0.009] (Fig.1b), comparable to effects of the same gene silencing approach in mature mice^17^. Note that incomplete protein loss was expected, given that tissue punches processed by western blot contained both transduced and unaffected tissues, and CaMKIIα-driven AAVs would be expected to spare glial β1-integrins, which are expressed at high levels^18^. (Western blot was used because immunostaining brain tissue with currently-available antibodies is quite difficult.) Figures depict group means and individual mice throughout.

Neuronal β1-integrin activation results in the phosphorylation of p190RhoGAP^19^. β1- integrin protein levels correlated with the proportion of phospho-p190RhoGAP/p190RhoGAP in knockdown mice, such that mice with fewer β1-integrins also had the less phospho-p190RhoGAP [*r*^*2*^=0.490, *p*=0.036] (Fig.1c), evidence that our gene silencing approach had functional consequences for integrin-mediated intracellular signaling.

The PL is necessary for mice and rats to use associations between actions and outcomes to guide choice behavior^13^. To understand whether β1-integrins in adolescence are necessary for this function in adulthood (*i*.*e*., to determine whether β1-integrins equip developing mice with the capacity for flexible choice behavior in adulthood), we reduced β1-integrins in adolescence and tested reward-related response strategies in adulthood. For comparison, we also reduced β1- integrin presence in adulthood. Mice were trained to nose poke at 2 apertures for 2 distinct food pellets (Fig.2a). Groups did not differ in response acquisition, with mice increasing responding over time [main effect of day (*F*_(6,312)_=33.26, *p*<0.0001), no day*group interaction (*F*_(18,312)_=1.037, *p*=0.4180), no other main effects (*F*s<1)] (Fig.2b). Further, there were no systematic preferences for either flavor of pellet that would affect subsequent phases of testing [*F*s<1; not shown], so response rates for both pellets are collapsed for simplicity.

**Figure 2.**
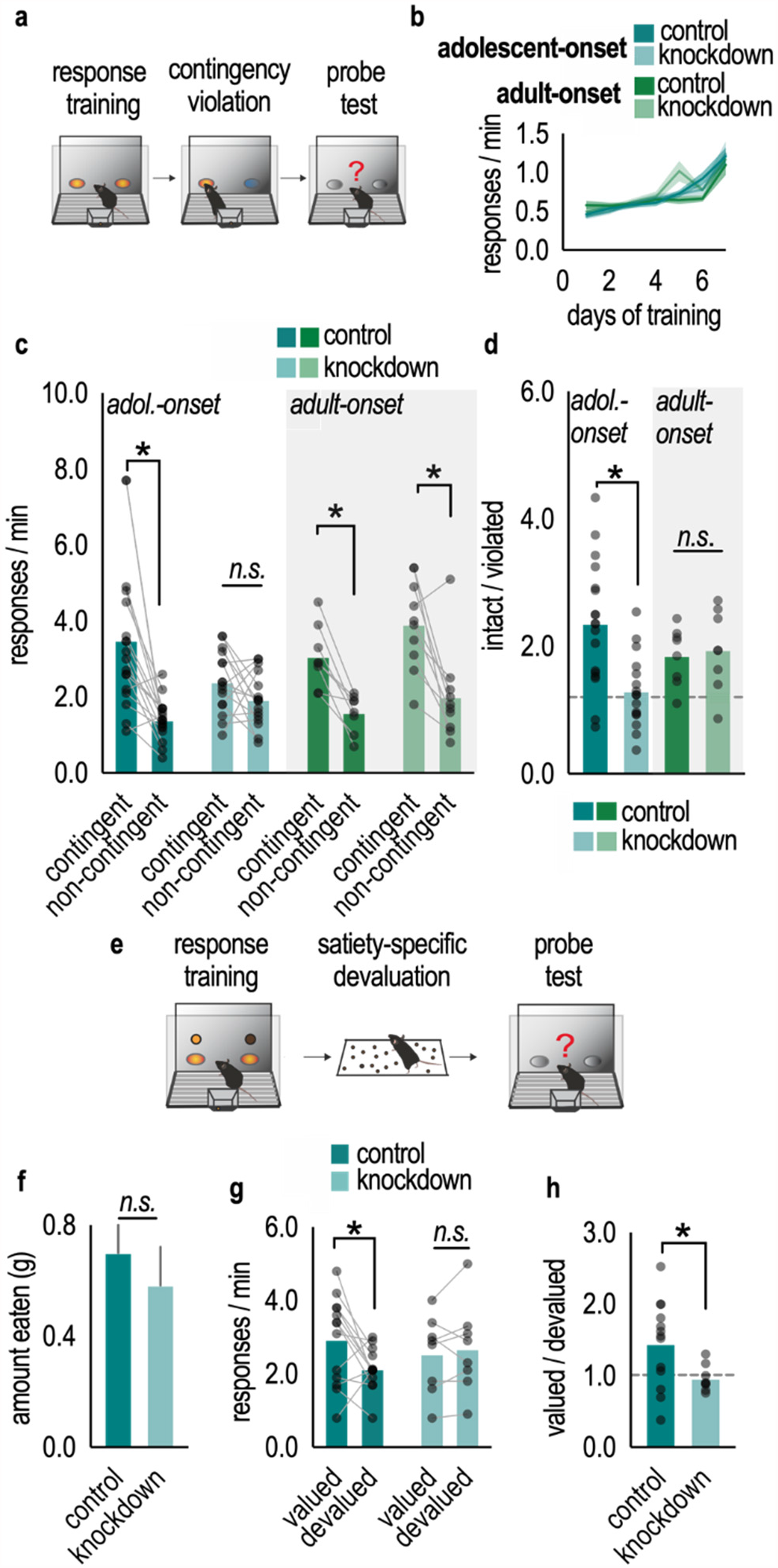
Developmentally-selective control of reward-related action by cell adhesion factor β1-integrin. **a**. Schematic. Mice are trained to nose poke at 2 apertures for 2 food reinforcers. Once mice are proficient, the causal relationship between 1 behavior and its outcome is violated. Preferential engagement of the behavior associated with the other, intact contingency is considered flexible action. **b**. Groups did not differ in response training. There were no systematic preferences for either flavor of pellet, so response rates for both pellets are collapsed here **c**. Control mice subsequently inhibited a behavior that was not reinforced and favored a reinforced behavior, but reduction of β1-integrins starting in adolescence ablated response preference. Adult-onset loss had no effects. **d**. The same data can be converted to response ratios (contingent/non-contingent condition). Loss of β1-integrins during adolescence, but not adulthood, reduced ratios, indicating that response performance is inflexible. **e**. Schematic. Mice are trained to nose poke at 2 nose poke apertures for 2 food reinforcers. Once proficient, mice are then allowed to freely consume 1 of the 2 food reinforcers in a separate environment. In a subsequent probe test, preferential responding for the other food (“valued” condition) is considered goal-directed. **f**. Groups did not differ in food consumed during the prefeeding session. **g**. Control mice preferred the aperture associated with the valued food reinforcer, whereas loss of β1-integrins ablated this preference. **h**. Accordingly, preference ratios (valued/devalued) were lower in integrin-deficient mice. Dashed lines at 1 indicate no preference. Bars represent means, error bars represent SEMs, and symbols represent individual mice. \**p*<0.05. *n*=8-18/group.

Next, the contingency between 1 familiar action and its outcome was violated by providing pellets non-contingently, or “for free.” In reaction, control mice inhibited that response during a probe test, preferring instead the response associated with the intact contingency. Meanwhile, loss of β1-integrins starting in adolescence, but not adulthood, ablated this preference [viral vector*age of knockdown*aperture interaction (*F*_(1,48)_=5.675, *p*=0.021), no other interactions (*F*s<1), main effect of aperture (*F*_(1,48)_=47.09, *p*<0.001), no main effect of viral vector (*F*<1) or age of knockdown (*F*_(1,48)_=2.12, *p*=0.151)] (Fig.2c).

We also compared response preference scores, calculated by dividing responses associated with the intact/violated contingencies. A ratio of >1 reflects a preference for the intact contingency – *i*.*e*., flexible action – while scores of 1 indicate no preference. The response ratio was ∼1 upon loss of β1-integrins during adolescence, while other groups generated higher scores [viral vector*age of knockdown interaction (*F*_(1,46)_=6.484, *p*=0.014), main effect of viral vector (*F*_(1,46)_=4.629, *p*=0.037), no main effect of age of knockdown (*F*<1)] (Fig.2d). Thus, developmental integrins are necessary for the ability of mature mice to engage in flexible reward-related action.

Goal seeking involves 2 processes: selecting behaviors based on causal knowledge – that is, knowledge that an action will lead to desired outcomes, as tested above – and selecting actions based on the value of likely outcomes^13^. Thus, we hypothesized that developmental β1-integrins would also be necessary for mice to select actions based on outcome value, which we tested using satiety-specific devaluation (Fig.2e). Given that β1-integrin loss in adulthood had no effects above, we focused on adolescent-onset β1-integrin reduction. Following training, mice were allowed to freely consume 1 of the 2 food reinforcers in a separate environment, decreasing the value of that reinforcer. Groups did not differ in food consumption [*t*_(14)_=0.6750, *p*=0.5107] (Fig.2f). Nevertheless, only control mice preferentially responded for the other, “valued” food when returned to the operant conditioning chambers, evidence that they selected actions based on outcome value. Meanwhile, reducing β1-integrins in adolescence blocked the ability of adult mice to select actions based on outcome value [viral vector*value interaction (*F*_(1,19)_=4.395, *p*=0.0497), no main effect of viral vector (*F*<1), no main effect of value (*F*_(1,19)_=2.531, *p*=0.154)] (Fig.2g). Accordingly, response ratios were lower upon loss of β1-integrins, reflecting no preference for the valued outcome [Welch’s corrected *t*_(15.40)_=2.660, *p*=0.0168] (Fig.2h). Thus, developmental integrins are necessary for mature mice to engage in goal-oriented action.

### BLA→PL projections are necessary for the expression of learned reward-related action

β1-integrins are largely localized in the post-synapse of excitatory synapses in the postnatal brain, coordinating the stability of these synapses^20, 21^. Thus, it seems likely that reducing *Itgb1* during adolescent development disrupts the ability of mature PL neurons to effectively receive inputs necessary for goal-seeking behavior. We next attempted to identify sources of such inputs. One candidate region is the BLA, given that BLA→PL projections develop throughout adolescence^22^; this development likely ensures that connections required for certain goal-seeking behaviors are accessible in adults^23, 24^. To determine whether these principles apply to the response strategies tested here, we first tested whether BLA→PL projections, independent of β1- integrins, are indeed necessary for action selection based on reward likelihood. We used a combinatorial viral vector strategy, infusing a retro-Cre into the PL and Cre-dependent Gi-coupled Designer Receptors Exclusively Activated by Designer Drugs (Gi-DREADDs+mCherry) into the BLA (Fig.3a). This combination would be expected to decrease the activity of BLA→PL projections in the presence of the DREADDs ligand, Clozapine N-oxide (CNO). Mice in the control group were infused with the same viral vector in the PL and a control viral vector lacking DREADDs in the BLA.

**Figure 3.**
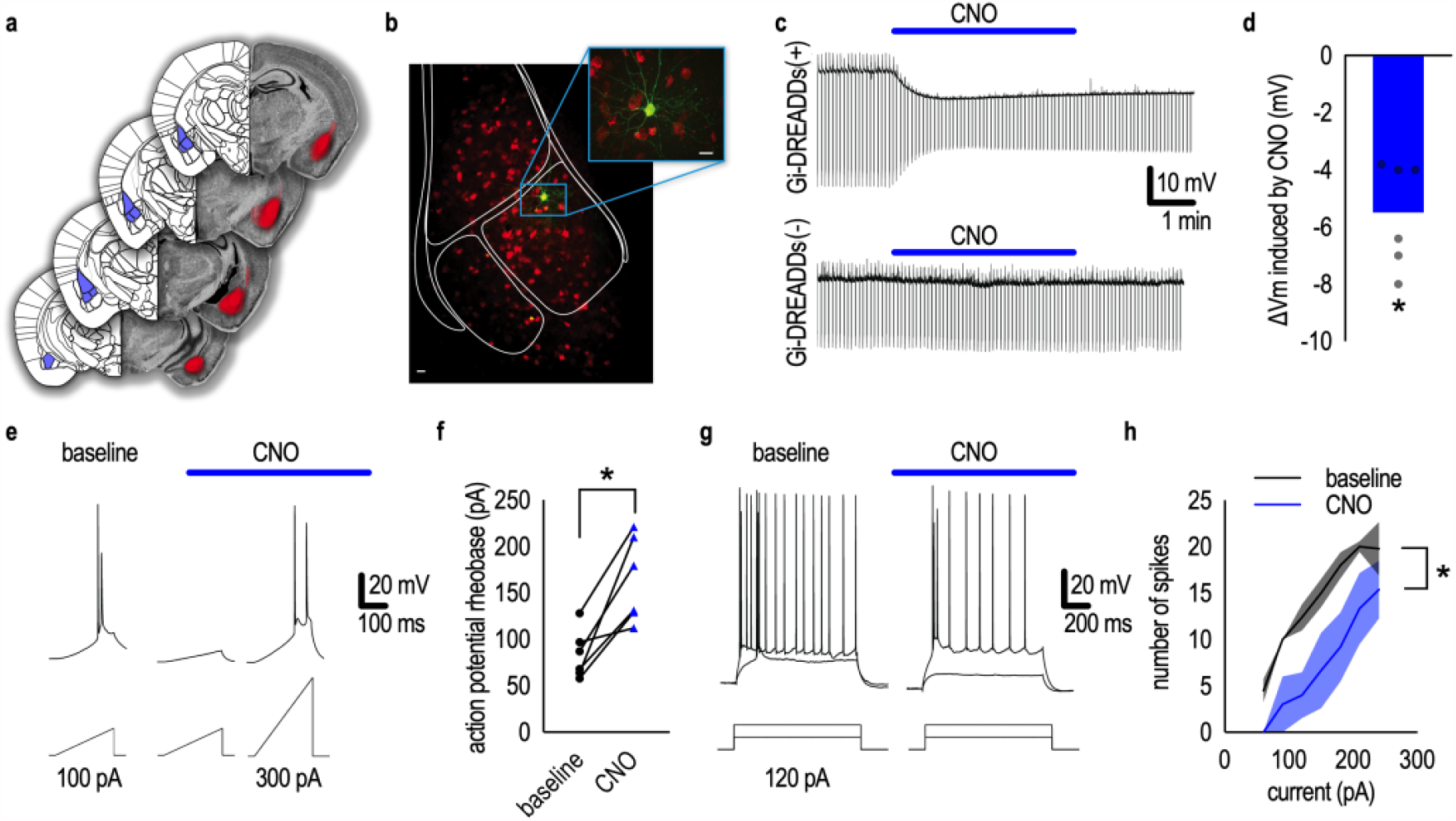
Chemogenetic silencing of PL-projecting BLA neurons. **a**. To inhibit the activity of projections from the BLA to the PL, mice were infused with viral vectors expressing a Cre-dependent Gi-DREADD into the BLA and a retrograde Cre into the PL. (**left**) Images from Allen Brain Atlas^70^ (from left: -1.28 mm, -1.64 mm, -2.12 mm, and -2.72 mm relative to bregma) depicting the BLA (purple). (**right**) Overlays of viral vector spread (red) encompassing the BLA on images from Mouse Brain Library^69^. **b**. Representative image of biocytin-filled (green) BLA neuron expressing Gi-DREADD-mCherry (red); Scale bar=20 µm. **c**. In a Gi-DREADD(+) neuron, bath application of CNO (10 μM) induced fast membrane hyperpolarization and decreased input resistance (upper trace). In a Gi-DREADD(-) neuron, CNO did not induce changes of membrane potential and input resistance (lower trace). **d**. CNO-induced hyperpolarization in Gi-DREADD(+) neurons. **e**. Rheobase, the minimal current required to generate action potential, was elevated upon CNO application. **f**. CNO increased rheobase in Gi-DREADD(+) neurons. **g**. CNO decreased the number of action potentials induced by 2 current steps. **h**. CNO decreased action potentials at different current steps. Bars represent means, error bars represent SEMs, and symbols represent individual neurons. \**p*<0.05. *n*=6 neurons from 4 mice.

To confirm the effectiveness of Gi-DREADDs, whole cell patch clamp recordings were obtained from BLA neurons (Fig.3b). In Gi-DREADDs-mCherry(+), but not Gi-DREADDs-mCherry(-), neurons, bath application of CNO (10 μM) induced rapid and sustaining membrane hyperpolarization [*t*_(5)_=7.418, *p*=0.0007] (Fig.3c,d). CNO also increased rheobase, or the minimum current required to generate a single action potential [paired *t*_(5)_=4.554, *p*=0.006] (Fig.3e,f). Lastly, we examined action potential firing induced by current injections. CNO reduced the number of spikes [main effect of CNO (*F*_(1,40)_=21.24, *p*<0.001), main effect of current (*F*_(6,40)_=9.049, *p*<0.001), no current*CNO interaction (*F*_(6,40)_=0.277, *p*=0.944)] (Fig.3g,h). In summary, Gi-DREADDs decreased the excitability of PL-projecting BLA neurons when CNO was administered, as expected.

Separate mice containing the same viral vectors (Fig.4a,b) were trained and then behaviorally tested. In the task in which we tested behavioral sensitivity to contingency, the memory encoding and expression periods occur on separate days. This, in combination with the ability to inducibly silence neurons in a temporally-specific fashion using DREADDs (unlike with *Itgb1* silencing), allows us to disentangle whether BLA→PL projections are involved in memory encoding or expression (Fig.4c). CNO was delivered to all mice either before a session when a familiar contingency was modified (during memory encoding), or ahead of the probe test (when mice must express memories in making choices). Mice were tested in both conditions in a counter-balanced fashion, and responses of control mice (mice bearing a control viral vector lacking DREADDs in the BLA) were averaged across both testing conditions.

**Figure 4.**
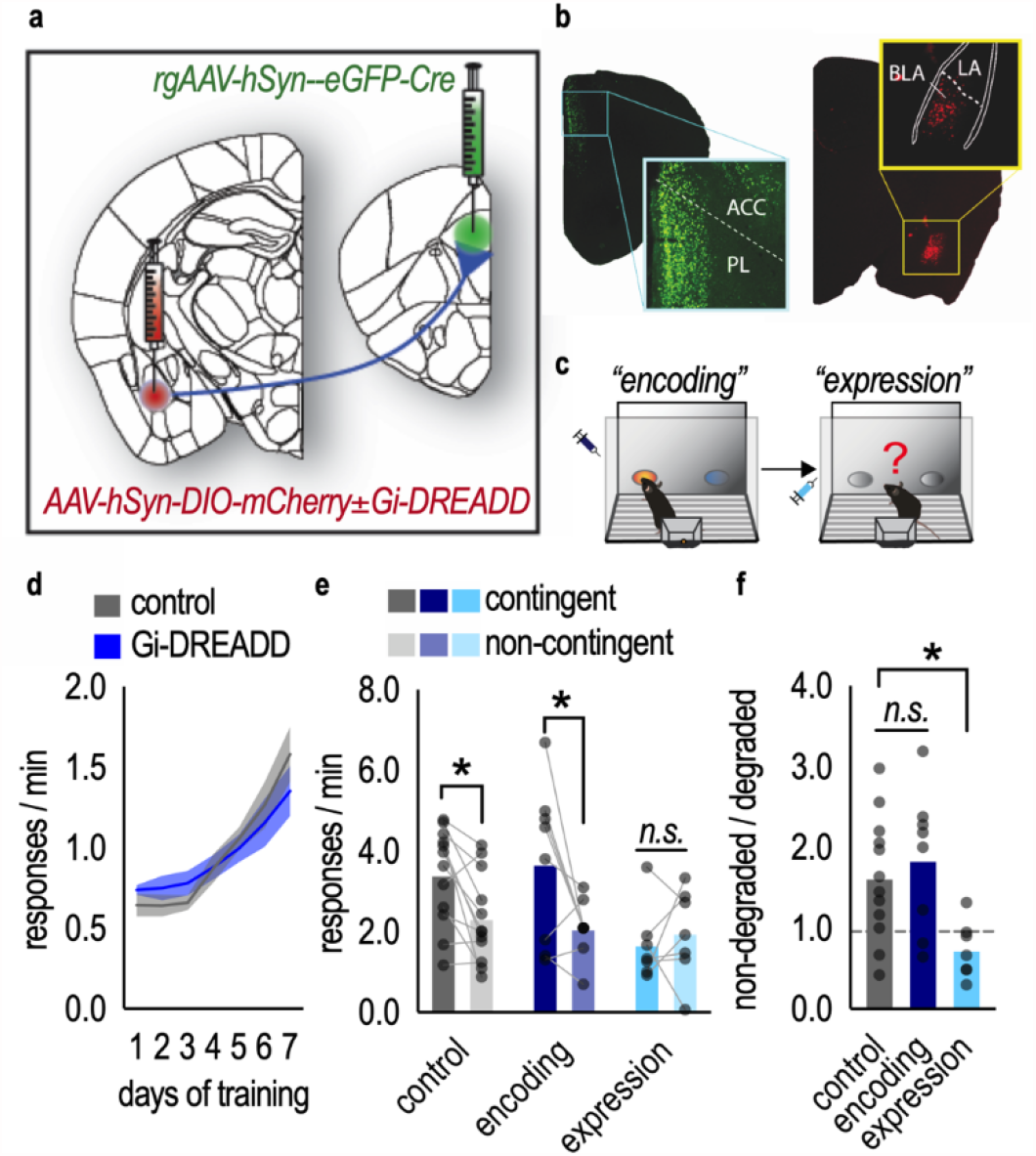
BLA→PL connections are necessary for flexible reward-related memory expression. **a**. Schematic of viral vector approach. Mice were infused with viral vectors expressing a Cre-dependent Gi-DREADD into the BLA and a retrograde Cre into the PL. Control mice received a Cre-dependent fluorophore, but no DREADD, in the BLA. **b**. Representative infusions in the BLA and PL. **c**. Mice received CNO either prior to the session when familiar action-outcome contingencies are being disrupted and new action-outcome memories are being **<u>encoded</u>**, or prior to the probe test, when mice are **<u>expressing</u>** recently formed action-outcome memories. **d**. Groups did not differ in response training. **e**. BLA→PL projection inactivation during the memory “encoding” period had no effect, while BLA→PL inactivation during the “expression” period ablated flexible response shifting. **f**. The same data are expressed as response preferences. Again, BLA→PL projection inactivation during the “encoding” period had no effect, while BLA→PL projection inactivation during the “expression” period ablated preference. Dashed line at 1 indicates no preference. Bars represent means, error bars represent SEMs, and symbols represent individual mice. \**p*<0.05. ACC=anterior cingulate cortex, LA=lateral amygdala. *n*=7-12/group.

Groups did not differ in response acquisition [main effect of day (*F*_(6,108)_=17.36, *p*<0.0001), no other main effects or interactions (*F*s<1)] (Fig.4d). Silencing BLA→PL projections during the probe test, but not earlier, ablated response preferences [group*aperture interaction (*F*_(2,24)_=3.424, *p*=0.042), main effect of aperture (*F*_(1,24)_=7.894, *p*=0.0097), no main effect of group (*F*_(2,24)_=2.884, *p*=0.0754)] (Fig.4e). Accordingly, response ratios were lowest following BLA→PL inactivation during the probe test [*F*_(2,24)_=5.904, *p*=0.008] (Fig.4f). These patterns suggest that BLA→PL connections are necessary for mice to express memories regarding reward likelihood and act accordingly, but not to encode those memories *per se*.

### Developmental β1-integrins are necessary for the morphological features of PL neurons positioned within a BLA-PL-striatal circuit

Our findings indicate that: **1)** developmental β1-integrins in the PL and **2)** BLA→PL connections are necessary for action flexibility in adulthood. These observations led to the hypothesis that β1-integrin presence sculpts the development of PL neurons receiving input from the BLA. Cortically-oriented BLA projections terminate on layer II, III, and V PL neurons^25^. We focus here on layer V neurons because these cells receive input from the BLA and send outputs to the dorsomedial striatum (DMS), which is required for the translation of decision-making strategies into action^26, 27^.

It was first necessary to isolate and characterize PL neurons receiving inputs from the BLA and projecting to the DMS under typical (β1-integrin+) conditions. We isolated projection-defined PL neurons by infusing an anterograde transsynaptic Cre-expressing viral vector into the BLA and a retrograde Cre-dependent mCherry-expressing viral vector into the DMS of adolescent *Thy1-*YFP^H^ mice^28^ (Fig.5a-b). Thus, mCherry+, YFP+ co-labeled cells represent layer V neurons that receive projections from the BLA, and project to the DMS (Fig.5c). For comparison, we infused an mCherry-expressing viral vector into the PL of adolescent and adult mice, allowing us to visualize a general population of transduced layer V neurons not defined by projection (Fig.5d). As above, viral vectors were infused at P21-24 or P56-60, and mice were euthanized 3 weeks later (Fig.5e). We then utilized high resolution imaging and 3D dendritic spine reconstruction to analyze the dendritic micro-architecture of each population.

**Figure 5.**
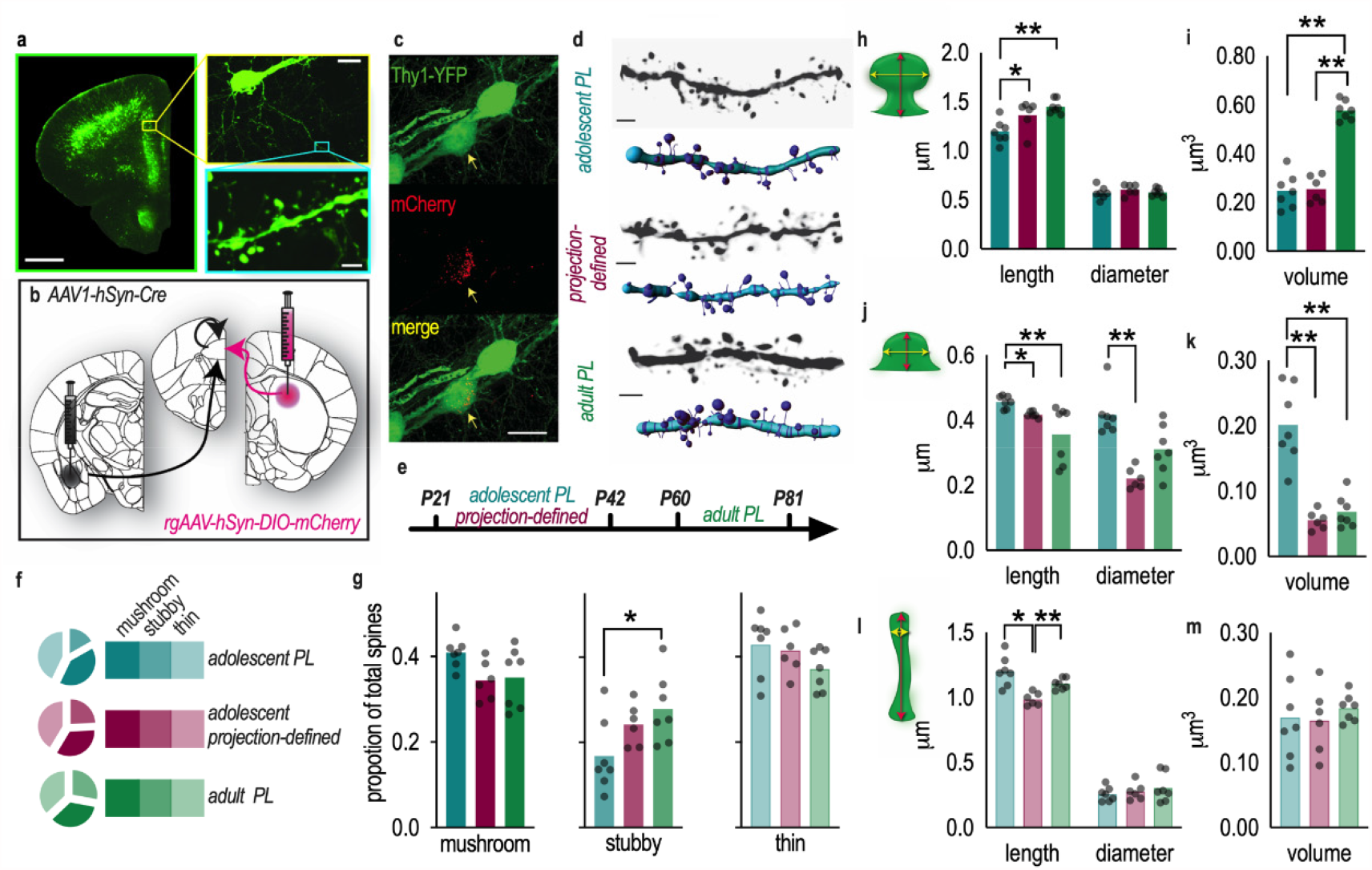
Morphological features of PL neurons receiving input from the BLA and projecting to the DMS. **a**. Coronal sections from YFP+ mice used to examine dendritic spine architecture on excitatory layer V PL neurons. Scale bar=1 mm (green), 20 μm (yellow), 2 μm (blue). **b**. Schematic of viral vector approach. A retrograde viral vector expressing a Cre-dependent fluorophore was infused into the DMS. A transsynaptic anterograde viral vector encoding Cre was infused into the BLA. YFP+mCherry+ PL neurons were imaged. This condition is referred to as “projection-defined.” Separate mice received an AAV-mCherry in the PL to transfect a general neuron population not defined by projection pattern. **c**. Representative PL neuron co-expressing YFP and mCherry (arrowhead). Scale bar=20 μm. **d**. Representative dendrites, with their associated 3D reconstructions. Scale bar=2 μm **e**. Timeline of experiments portraying time of viral vector delivery (tick mark at the left of text) and euthanasia (tick mark at the right of text), with the three conditions labeled in different colors. **f**. Pie charts representing the dendritic spine subtype proportions in each group. **g**. Mature neurons had the highest proportion of stubby-type spines, while projection-defined neurons exhibited an intermediate phenotype. There were no significant differences in the proportions of mature- or thin-type spines. **h**. Projection-defined adolescent PL neurons had longer mushroom-type spines than the general adolescent population, resembling mature morphologies. We observed no differences in dendritic spine diameters. **i**. Mature neurons had mushroom-shaped spines of the largest volume, exceeding both adolescent populations. **j**. Projection-defined adolescent PL neurons and mature PL neurons had shorter stubby-type spines than the general population of adolescent PL neurons. Projection-defined adolescent PL neurons and mature neurons had smaller stubby-type spine diameters than the general adolescent population. **k**. Their volumes were also lower. **l**. Projection-defined adolescent PL neurons had shorter thin-type spines than other groups, and there were no differences in thin spine diameters, or **m**. thin spine volumes. Bars represent means, and symbols represent individual mice. \**p*<0.05, \*\**p*<0.001. *n*=6-7/group.

Interestingly, projection-defined PL neurons had lower dendritic spine densities than either adult or adolescent undefined PL layer V populations (fig.S1). To account for these differences, we focused here on the proportions of mature and immature spine subtypes, given that the balance between mature *vs*. immature spine subtypes allows for the prioritization and integration of certain inputs over others^29^. The proportion of immature, stubby-shaped dendritic spines was highest in adulthood and lowest in adolescence, and projection-defined neurons had an intermediate phenotype, indistinguishable from both groups [*F*_(2,17)_=3.914, *p*=0.04] (Fig.5f,g). Projection-defined neurons had a lower proportion of mature, mushroom-shaped dendritic spines than the general population of adolescent neurons, more closely resembling adult neurons, although this comparison did not reach statistical significance [*F*_(2,17)_=3.115, *p*=0.0704] (Fig.5f,g). Thin-type spine proportions did not differ between groups [*F*_(2,1\7)_=1.806, *p*=0.1945] (Fig.5f,g).

We next characterized the morphological features of each dendritic spine subtype. Mushroom-type spines on projection-defined neurons again more closely resembled adult spines in length than same-age counterparts collected from a general layer V population [*F*_(2,17)_=8.578, *p*=0.0027] (Fig.5h). Spine diameters did not differ [*F*<1] (Fig.5h). Interestingly, spine volume was largest in adult mice [*F*_(2,17)_=74.37, *p*<0.0001] (Fig.5i). Thus, the morphology of mushroom-shaped spines on projection-defined neurons in some ways resembled adult spines (in length), and in some ways resembled their undefined adolescent counterparts (in volume).

We identified a similar pattern for stubby-shaped spines, in that stubby-type spines on projection-defined neurons exhibited decreased length [*F*_(2,17)_=6.036, *p*=0.0105] and diameter [*F*_(2,17)_=15.94, *p*=0.0001] compared to the general population of same-age spines and more closely resembled adult spines (Fig.5j). Stubby-type spine volume was similar in adult and projection-defined adolescent populations, and far lower than a general adolescent population [Welch’s ANOVA *W*_(2,10.19)_=18.76, *p*<0.0004] (Fig.5k).

Lastly, projection-defined adolescent neurons had shorter thin-type spines than other groups [*F*_(2,17)_=12.28, *p*=0.0005], with no group differences in diameters or volumes [*F*s<1] (Fig.5l,m). Thus, layer V PL neurons positioned within a BLA→PL→DMS circuit exhibit distinctive morphologies relative to a general population of layer V PL neurons.

We next asked: Do β1-integrins stabilize neuron structure in a BLA→PL→DMS circuit? *And in particular, during an adolescent sensitive period*? To address these questions, we felt it important to integrate into our experimental design a condition in which mice lost β1-integrins *prior to* adolescence, to complement conditions in which β1-integrins were lost *during* or *following* adolescence. Reducing β1-integrins prior to adolescence via viral vector delivery is difficult due to the small size of neonatal mice, so we instead crossed YFP+ *Itgb1*-flox mice with mice expressing *Emx1-*Cre to achieve forebrain-specific loss of *Itgb1* starting at embryonic day 11.5, when the *Emx1* promoter becomes active^30^. Other groups of *Itgb1*-flox mice received either control or Cre-expressing viral vectors as in prior figures and summarized in table 1. Therefore, we were able to compare the effects of embryonic-, adolescent-, and adult-onset β1-integrin loss on layer V PL neurons, including those positioned in a BLA→PL→DMS circuit. A timeline reflecting gene silencing and euthanasia timing is provided in Fig.6a.

Dendritic spine imaging and reconstruction revealed that β1-integrins in adolescence, but not earlier or later, control dendritic spine density (fig.S1) and subtype distribution on PL neurons. Proportions of mushroom- and thin-type spines were lower in both adolescent β1-integrin-deficient conditions (*i*.*e*., general layer V neurons and those positioned in a BLA→PL→DMS circuit), and comparisons were statistically significant on projection-defined neurons [projection-defined mushroom (*t*_(11)_=2.400, *p*=0.0373), thin (*t*_(11)_=2.235, *p*=0.0452); general population mushroom (*t*_(12)_=1.972, *p*=0.0721), thin (*t*_(12)_=1.948, *p*=0.0752)] (Fig.6b,c). β1-integrin reduction in adolescence accordingly elevated proportions of stubby-type spines, as expected [projection-defined (*t*_(11)_=6.758, *p*<0.0001); general population (*t*_(12)_=2.353, *p*=0.0365)] (Fig.6b,c). When we calculated the proportion of mushroom to non-mushroom spines, projection-defined PL neurons again suffered a significant loss [*t*_(11)_=2.745, *p*=0.0191], reflecting lower proportions of mature spines – those likely to contain synapses (Fig.6d).

In contrast to the adolescent groups, embryonic-onset and adult-onset β1-integrin reduction had no effects [*p*>0.05] (fig. 6b,c), even despite gross cortical layering deficits following embryonic gene silencing. This phenomenon was first reported by ref.^31^ and is replicated in fig.S2.

**Figure 6.**
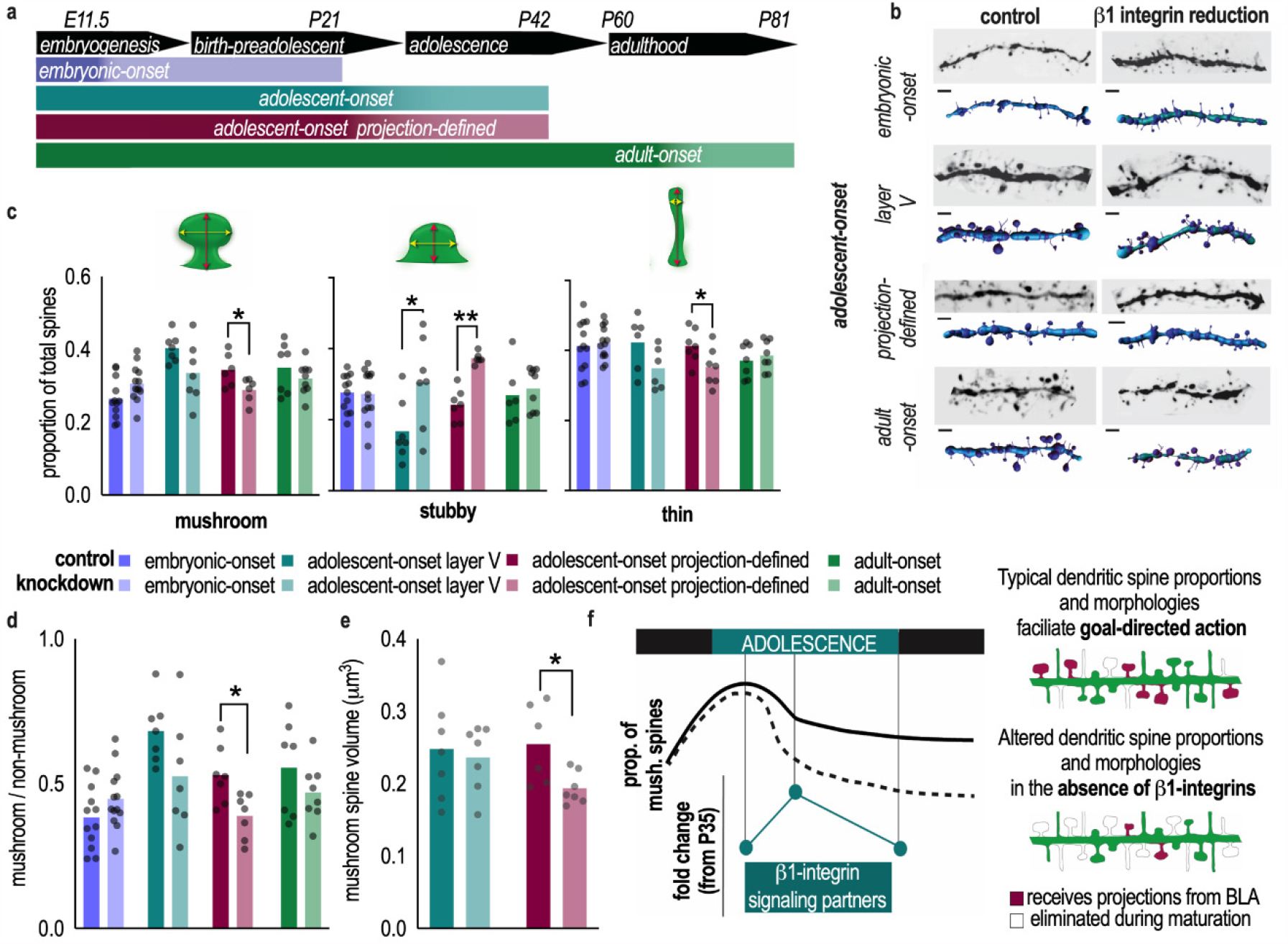
Cell adhesion control of PL neuronal morphology. **a**. Timeline. β1-integrins were reduced starting in embryogenesis, adolescence (globally or specifically in projection-defined BLA→PL→DMS neurons), or adulthood. Timing of diminishing β1-integrin tone is represented by thinning color gradient. Dendritic spine analysis occurred at P21, P42, or P81 for embryonic-, adolescent-, or adult-onset investigations, respectively. **b**. Representative dendrites, with their associated 3D reconstructions. Scale bar=2 μm. **c**. Adolescent-onset β1-integrin reduction decreased the proportion of mushroom- and thin-type dendritic spines and increased the proportion of stubby-type spines. Embryonic- and adult-onset β1-integrin reduction did not affect dendritic spine proportions. **d**. Adolescent-onset β1-integrin loss in projection-defined neurons decreased the proportion of mature mushroom-shaped spines to non-mushroom spines. **e**. β1-integrin tone during adolescence is also required for intact mushroom spine volume on projection-defined neurons. **f**. Model. Typical dendritic spine densities increase in the PL in the pre-adolescent period, then some dendritic spines are eliminated, and others are stabilized during adolescence. β1-integrin signaling partners increase in expression during adolescence^12^. Here, we show that β1- integrin presence stabilizes dendritic spines on PL neurons, including neurons that receive input from the BLA and project to the DMS. Absent this activity, typical proportions and morphologies of mature, mushroom-shaped spines are lost (dashed line), and PL function is compromised in adulthood. Bars represent means. Symbols represent individual mice. \**p*<0.05, \*\**p*<0.001. *n*=7- 13/group.

Finally, we compared the volumes of dendritic spines in our adolescent groups. In addition to changing spine proportions, β1-integrin loss decreased the volumes of mature, mushroom-shaped spines on projection-defined neurons [Welch’s corrected *t*_(6.408)_=2.535, *p*=0.0419], but not in the general population [*t*_(12)_=0.3584, *p*=0.7262](Fig.6e). Thus, β1-integrins sustain typical proportions and morphologies of mature, mushroom-shaped dendritic spines – those most likely to contain synapses – within a developing BLA→PL→DMS pathway (model in Fig.6f). We speculate that this function enables mature PL neurons to house stable synaptic connections necessary for goal-directed action in adulthood.

## Discussion

Across mammalian species, PFC development extends well into adolescence. Concurrent processes – dendritic spine elimination and stabilization – determine which inputs will be maintained until adulthood^32^. We report that the cellular adhesion factor β1-integrin in the PL subregion of the PFC stabilizes dendritic spines on excitatory neurons during adolescent development, and adolescent β1-integrin presence is required for PL function in adulthood, specifically in choosing actions based on their consequences. We find that the expression of learned reward-related action strategies requires BLA inputs to the PL. This pattern led us to then confirm that β1-integrins on developing PL neurons receiving projections from the BLA and projecting to the DMS, a primary striatal output controlling reward-related action^27^, are required for healthy dendritic spine micro-architecture later in life. We envision that β1-integrin-mediated dendritic spine stabilization during adolescence provides structural substrates for synaptic connections necessary for flexible action in adulthood.

### Cell adhesion during adolescence is required for flexible action later in life

Both humans and rodents are sensitive to the relationships between actions and their outcomes. Typical mice will change behaviors if causal relationships deteriorate, or if the value of a given outcome declines. We report that β1-integrins in the PL during adolescence are necessary for these action strategies to manifest later in life. Specifically, reducing β1-integrins in adolescence rendered mice less likely to select actions based on outcome likelihood or value. Lesions and inducible inactivation of the PL have the same effects^13, 14^, but the current report is the first, to the best of our knowledge, to establish the necessity of a cell adhesion receptor in these functions. Reducing β1-integrins in adulthood had no effects; thus, it appears likely that β1-integrins support goal-directed action via the stabilization of dendritic spines during adolescence, as opposed to any acute function in synaptic plasticity in adulthood. Importantly, these findings in mature mice do not negate the established function of the PL in flexible action selection in mature rodents; rather, they rule out β1-integrin as a factor that controls this capacity in the post-adolescent period.

### Flexible action requires BLA→PL connections

We found that silencing BLA→PL projections prohibited behavioral flexibility when mice had to retrieve a newly formed outcome-related memory to guide choice behavior. Generally, these findings are consistent with reports that pathway-specific ablation of BLA→PL projections renders mice insensitive to outcome value^24^ (but see^33^), indicating their involvement in goal-directed action. However, the PL is often thought to encode, but not necessarily retrieve, outcome-related memories (reviewed in^13^), so our finding was in other ways unexpected. In seminal studies, lesions were placed in the PL prior to instrumental training, or prior to any testing of goal-directed action^34-36^. This experimental design makes it difficult to disambiguate PL function(s) during different phases of training and testing. More recent experiments used inducible inactivation techniques to test PL function at different phases: during instrumental response training^37^, during the consolidation of new memory^38^, and during the expression of that memory^39^. Shipman and colleagues demonstrated that inducible inactivation of the PL impairs the retrieval of action-outcome memories following brief but not extended training^39^. PL neurons may thus form an “early” memory engram for outcome-related information as animals gain initial experience with a task, as has been reported in fear conditioning contexts^40^. Meanwhile, the BLA represents specific outcome information that is motivationally significant and necessary for choosing between options^23^. The contingency violation in our task causes a transient “extinction burst,” evidence of motivational salience^41^, so involvement of the BLA in subsequent response prioritization is perhaps unsurprising. Our findings indicate that BLA inputs to the PL help animals prioritize certain reward-seeking behaviors over others during memory retrieval epochs.

### Developmental β1-integrins are necessary for the morphological features of PL neurons positioned in a BLA→PL→DMS circuit

The ability to select actions based on their consequences requires BLA projections to the PL (Fig.4) and PL outputs to the DMS^26, 27^. Further, the ability of PL→DMS projections to exert control over goal-directed action^26^ is likely subject to input from the BLA, as inactivation of the BLA prevents learning-related plasticity at PL→DMS synapses^24^. We thus visualized developing adolescent PL neurons positioned within this di-synaptic circuit, receiving projections from the BLA and projecting to the DMS. We enumerated: thin-type dendritic spines, which exhibit rapid turnover with a high potential for involvement in learning and memory; stubby-type dendritic spines, a transitory phenotype that does not contain synapses; and mushroom-shaped spines, which house high densities of glutamate receptors and stable synapses^42^. In some instances, these projection-defined adolescent PL neurons more closely resembled mature PL neurons than a general population of same-age adolescent neurons, for instance in the proportion of mature spines detectable on dendrites.

One notable difference was that the mushroom-shaped spines on adult neurons were larger in volume than those on adolescent neurons, including projection-defined adolescent neurons. Larger spines are less motile and less plastic than other spines^29^, containing large stable synapses^43^ and a high density of AMPA receptors^44, 45^. In contrast, smaller dendritic spines contain relatively more NMDA receptors^46^, are highly sensitive to Ca^2+-^mediated signaling^47^, and exhibit a greater capacity for activity-dependent synaptic strengthening. Thus, adolescent neurons appear to be primed for new learning, with high densities of small mushroom-shaped spines. Reinforcement learning and the capacity for action selection according to action-outcome contingencies improve throughout adolescence^48, 49^. Likely mechanistic factors driving this improvement include the strengthening of BLA→PL connectivity^22^, the elimination of extraneous dendritic spines on PL neurons^12, 50^, and as we reveal, the integrin-mediated maturation of dendritic spines on PL neurons receiving input from the BLA and projecting to the DMS (Fig.5).

To determine whether β1-integrin presence controls PL neuron development during an adolescent sensitive period, we integrated into our anatomical investigation a group of mice with *embryonic*-onset gene silencing, to complement neurons that had undergone *adolescent*- or *adult*-onset gene silencing. Despite the inclusion of these multiple conditions, β1-integrin loss only in adolescence detectably modified dendritic spine subtype presence, biasing dendritic spine subtypes towards stubby-type spines at the expense of mushroom- and thin-type spines (those with higher potential for synaptic presence). This effect was most profound in projection-defined neurons. Lower-than-typical proportions of mature spine types likely imperils signal fidelity from incoming projections. We speculate that β1-integrin tone in adolescence confers structural stability, preserving synaptic efficacy^44, 45^. Supporting this perspective, β1-integrins signal in neurons through p190RhoGAP and ROCK2 to encourage actin filament proliferation^51^, thereby maintaining dendritic complexity^10^, and prefrontal cortical levels of these proteins peak in adolescence^12^. Further, β1-integrins promote interactions between the Arp2/3 complex and cortactin, supporting actin branching and polymerization^52-54^, processes that would enable the maturation of spines from an immature to mushroom morphology capable of housing synapses. Branching and actin polymerization would also counter the retraction or elimination of mushroom-type spines. *In vivo* imaging could disentangle precise β1-integrin functions in developing neurons at far more granular levels (*e*.*g*., see^50^).

Notably, β1-integrins are involved in activity-dependent plasticity: For instance, integrin ligands do not alter baseline physiological properties of cultured hippocampal neurons, but they stabilize long-term potentiation (LTP)^55^. Further, integrin blocking antisera disrupts LTP-induced actin polymerization and consolidation^11,56^. Multiple aspects of dendritic spine maturation in adolescence are activity-dependent^32^, thus the recruitment of β1-integrins seems sensible. Meanwhile, loss of β1-integrins *before* adolescence here had no effects on dendritic micro-architecture. Similarly, β1-integrins in embryogenesis do not appear to impact dendritic spine densities on hippocampal CA1 neurons, even while they are involved in the proper maturation of presynaptic physiology^31^, likely via the signaling partner talin^57^. We also found no effects of β1- integrin reduction in adulthood. This outcome might highlight the exquisite capacity of cell adhesion factors to compensate for one another in mature organisms^21^.

## Conclusions

To summarize, we find that presence of the cell adhesion molecule β1-integrin during adolescence is essential for flexible action later in life. We attribute this function to β1-integrin-mediated control dendritic spine restructuring that occurs during adolescent development, for instance, stabilizing PL neurons receiving input from the BLA. These neurons are necessary for retrieving memories linking actions and outcomes. Thus, it seems likely that destabilizing their capacity to house stable synapses (via developmental β1-integrin loss) imperils the capacity of mice to select actions based on likely outcomes.

Integrin signaling is thought to be dysregulated in substance use disorders^58^ and schizophrenia^59^, diseases that often form or first present in adolescence and in which defects in the capacity for goal-directed action are hallmark. Several β1-integrin signaling partners are pharmacologically modulable, opening avenues for new clinical applications. For instance, stimulating β1-integrin-mediated signaling pathways during adolescence could potentially combat decision-making difficulties that are characteristic, and often reinforcing, of illness^60^. More broadly, better understanding which cell adhesion molecules support adolescent brain development (and how) may prove to be fertile ground in developing new therapeutic approaches.

## Materials and Methods

### Subjects

Experiments used transgenic *Itbg1*^*tm1Efu*^ (*Itgb1-*flox) mice bred on a mixed strain background (C57BL/6J;129×1/SvJ). These mice have loxP sites flanking exon 3 of *Itgb1*; Cre introduction deletes this exon^16^. Control mice were littermates that received a control viral vector, rather than a Cre-expressing viral vector. *Itgb1-*flox mice were crossed with mice expressing YFP under the *Thy1* gene (H line^28^) for dendritic spine imaging. Again, control mice were littermates that received a control viral vector, rather than a Cre-expressing viral vector, or littermates lacking *Itgb1*-flox (see table 1 for conditions for each experiment). For embryonic-onset *Itgb1* silencing, *Itgb1-*flox/*Thy1-*YFP mice were crossed with mice expressing *Emx1*-Cre, allowing for forebrain-specific expression of Cre beginning at embryonic day (E) 11.5^30^. Control mice in this experiment were *Thy1-*YFP/Cre+ littermates lacking *Itgb1-*flox. In this condition, *Itgb1* would be unaffected. For projection-specific inactivation studies, C57BL/6J mice bred in-house from Jackson Labs stock were used, and viral vector conditions and controls are outlined in table 1. Throughout, mice were randomized to group.

Table 1 summarizes which mice were used in each experiment and ages of manipulations. For *Itgb1* knockdown experiments, mice infused with Cre-expressing viral vectors between postnatal days 21-24 are referred to as the “adolescent-onset” group because viral-mediated gene silencing would be expected to occur during adolescence, generally thought to start at postnatal day 28 in rodents. Meanwhile, Cre-expressing viral vectors were delivered at or soon after P56 in the adult condition to achieve gene silencing in adulthood.

Throughout, both sexes were used, and we did not detect sex differences. Mice were maintained on a 12 hr light cycle (0700 on) and provided food and water *ad libitum* unless otherwise noted. Procedures were approved by the Emory University IACUC.

### Stereotaxic surgery and viral vectors

Mice were anesthetized with ketamine/dexmedetomidine (100 mg/kg/0.5 mg/kg, intraperitoneal injection (*i*.*p*.)) and placed in a digitized stereotaxic frame (Stoelting). Small holes were drilled in the skull, and viral vectors were infused at AP+1.7mm, ML±0.17, DV-2.5 (PL); AP-1.4, ML±3.0, DV-4.9 (BLA); or AP+0.5, ML±1.5, DV-4.25 (DMS). Viral vectors were infused over 5 min in a volume of 0.5 μl. Viral vectors were supplied by UNC Viral Vector Core or Addgene and are described in table 1. Syringes were left in place for ≥5 min for PL infusions, or ≥8 min for BLA/DMS infusions, prior to removal and suturing. Mice were revived with antisedan (3 mg/kg, *i*.*p*.) and left undisturbed for at least 3 weeks prior to euthanasia (for immunoblotting or dendritic spine analysis) or behavioral experiments.

### Immunoblotting

Mice were euthanized via rapid decapitation after brief anesthesia with isoflurane. Brains were extracted, frozen at -80 °C, and sectioned using a chilled brain matrix into 1 mm slices. Tissue punches containing the PL were extracted using a 1 mm diameter tissue core. Tissues were homogenized in lysis buffer ((200 µl: 137 mM NaCl, 20 mM tris-HCl (pH=8), 1% igepal, 10% glycerol, 1:100 Phosphatase Inhibitor Cocktails 2 and 3 (Sigma), 1:1000 Protease Inhibitor Cocktail (Sigma)), and stored at -80°C. Protein concentrations were determined using a Bradford colorimetric assay kit (Pierce).

15 µg of protein was separated by SDS-PAGE on 7.5% gradient tris-glycine gels (Bio-Rad). Following transfer to PVDF membrane, blots were blocked with 5% nonfat milk for 1 hr. Membranes were subsequently incubated with primary antibodies: β1-integrin (rabbit, Cell Signaling, 4706, 1:200), anti-RhoGAP p190 (mouse, Millipore, 05378, 1:1000), anti-phosphotyrosine (mouse, generously provided by A. Koleske, clone 4G10, 1:500). p190RhoGAP is the predominant phosphotyrosine-containing protein of 190 kD recognized by the 4G10 antibody in mouse brain tissue^61^ so we used the phosphotyrosine antibody to detect phosphorylated p190RhoGAP.

After exposure to primary antibodies, membranes were incubated in horseradish peroxidase secondary antibodies (anti-mouse, 1:10,000, Jackson ImmunoResearch; anti-rabbit, 1:10,000, Vector Labs) for 1 hr. Immunoreactivity was assessed using a chemiluminescence substrate (Pierce) and measured using a ChemiDoc MP Imaging System (Bio-Rad). For β1- integrins, densitometry values were first normalized to a loading control, then normalized to the control sample mean from the same membrane in order to control for variance in fluorescence between gels. Phosphorylated p190RhoGAP levels were normalized to total p190RhoGAP. Immunoblots were replicated at least twice with concordant results, and proteins were detected at expected molecular weights.

### Instrumental response training

All mice were tested in adulthood (>P56). First, mice were food restricted to ∼90% of their free-feeding body weight to motivate food-reinforced responding. Mice were then trained to nose poke for 2 distinct food reinforcers (20 mg Bio-Serv Dustless Precision Pellets, grain and chocolate) in Med Associates operant conditioning chambers equipped with 2 nose poke apertures and a separate food magazine. Responding on each aperture was reinforced using a fixed ratio 1 (FR1) schedule of reinforcement, such that 30 pellets were available for responding on each aperture. Sessions ended at 70 min or 60 pellets acquired. Training ended after at least 7 sessions, when mice acquired all 60 pellets within 70 min. Response acquisition curves represent responses/min for both nose poke apertures during the last 7 training sessions, with no side or pellet preferences throughout.

### Test for action-outcome response selection

was conducted as previously described^17, 62^. First, 1 of the 2 apertures was available, and responding continued to be reinforced according to an FR1 schedule for 25 min. As such, the action-outcome contingency associated with that aperture remained intact. The following day, the other aperture was available for a 25 min session during which the action-outcome association between that nose poke behavior and reward were violated. In this case, pellets were delivered non-contingently at a rate matched to each animal’s reinforcement rate from the previous session. Responding was not reinforced. Thus, this nose poke action becomes significantly less predictive of reinforcement than the other^63^. The apertures, as well as the order of the “contingent” and “non-contingent” sessions, were counterbalanced.

Finally, both apertures were made available during a 15 min probe test conducted in extinction the following day. Preferential engagement of the response that remained reinforced is evidence of using contingency knowledge to guide choice. Meanwhile, engaging both responses equivalently reflects a failure in response updating^14^.

### Outcome devaluation

A version of classical instrumental outcome devaluation was used, as previously described^64^. Following response training, mice were placed into a clean, novel chamber with 1-2 g of either grain- or chocolate-flavored pellets for 30 min (males) or 90 min (females), devaluing that pellet. (In our experience, females require longer to consume this amount, hence the longer prefeeding period.) Immediately following, mice were placed in the operant conditioning chambers, and both apertures were available for a 15 min probe test conducted in extinction. Action selection according to outcome value is reflected by preferential responding for the valued relative to devalued pellet. If mice ate ≤0.2 g during the prefeeding period, the procedure was repeated. Due to technical error, pellet consumption of only 16/21 mice was recorded.

### Clozapine-N-Oxide (CNO) administration *in vivo*

In experiments using DREADDs, all mice received CNO (Sigma; 1 mg/kg, *i*.*p*., in 2% DMSO and saline), regardless of viral vector condition, to equally expose animals to any unintended consequences of CNO^65^. For projection-specific inactivation studies, CNO was administered 30 min prior to either the “non-contingent” session or the probe test. The procedure was then repeated, with mice receiving CNO at the opposite time point. The previously intact contingency was violated and *vice versa*. The control group is reported as an average response rate for each control mouse across both injection time points.

### Histology

For analysis of viral vector spread, mice were transcardially perfused under deep anesthesia with ketamine/xylazine (120 mg/kg/10 mg/kg, *i*.*p*.), prior to brain incubation in 4% paraformaldehyde. Brains were next transferred to 30% w/v sucrose solution, then were sectioned on a Leica microtome held at -15°C into 50 µm sections, mounted, and coverslipped. The mCherry tag was visualized. Immunohistochemistry for mCherry (mouse, 1:1000, Takara; goat anti-mouse-alexa594, 1:500, Invitrogen) was used to delineate viral vector spread, as needed.

### Electrophysiology

Mice were euthanized by rapid decapitation after brief anesthesia with isoflurane. Brains were extracted and sectioned using a vibrating microtome (Leica) into 300 µm sections. Sections containing the BLA were incubated in 95%O_2_/5%CO_2_ oxygenated 32°C sectioning ACSF for 1 hr before recording (for details, see^66^).

Brain sections were transferred to a recording chamber mounted on the stage of a Leica microscope and perfused with recording ACSF at a rate of ∼2 ml/min. BLA→PL projection neurons were identified by the expression of mCherry using a Leica STP6000 epifluorescence microscope. Whole cell patch clamp recordings were performed using a MultiClamp 700B amplifier, a Digidata 1550 digitizer and pClamp 10.6 software (Molecular Devices). Recording pipettes have resistance of 4∼6 MΩ when filled with potassium gluconate-based patch solution (in mM): K-Gluconate (130), KCl (2), HEPES (10), MgCl_2_ (3), phosphocreatine (5), K-ATP (2), and NaGTP (0.2). The patch solution was adjusted to pH 7.3 and had an osmolarity of 280-290 mOsm. 0.3% biocytin was included in the patch solution to stain and localize patched BLA neurons. Membrane potential was current clamped at -60 mV for baseline recordings and input resistance was monitored by observing membrane responses to hyperpolarization current injections (100-200 pA, 0.5 s).

To examine the inhibitory effects of Gi-DREADD activation on BLA neurons, we recorded 3 measures before and during CNO (10 µM) application: **1)** membrane potential hyperpolarization; **2)** rheobase, the minimum current required to generate a single action potential in a ramp current injection protocol (250 ms); and **3)** the number of spikes in response to current injection steps (1 s duration, delta 30-60 pA). Whole-cell access resistances were deemed acceptable if current clamp measurements were <20 MΩ and changed <15% throughout recording. Analysis was performed offline using Clampfit 10.6 (Molecular Devices).

### Dendritic spine imaging

Mice were euthanized by rapid decapitation after brief anesthesia with isoflurane, and brains were extracted and submerged in 4% paraformaldehyde. Brains were next transferred to 30% w/v sucrose solution, then were sectioned on a Leica microtome held at -15°C into 50 µm sections, mounted, and coverslipped. Unobstructed dendritic segments co-expressing mCherry and YFP (or just YFP in *Emx* mutant mice) were imaged on a spinning disk confocal (VisiTech International) on a Leica DM 5500B microscope. Z-stacks were collected with a 100x 1.4NA objective using a 0.1 μm step size, sampling above and below the dendrite. After imaging, we confirmed at 10x that the image was collected from the PL. For each animal, 5-12 independent basal dendrite segments, located 25-150 µm from the soma, were imaged.

Dendritic spines were reconstructed in 3-D using Imaris software and previously described methods^67^. Images were not manipulated prior to Imaris processing. Each experiment was scored by a single blinded rater. Each protrusion ≤4.5 µm in length was considered a spine. Bifurcated spines were considered singular units. Dendritic spines less than 0.6 µm in length were defined as stubby. If spine length was greater than 0.6 µm, spine classification was dependent on head diameter: spines with a head diameter more than 2 times the neck diameter were classified as mushroom, and spines with a head diameter less than 2 times the neck diameter were classified as thin^68^.

### Statistics

All statistics were performed using GraphPad Prism or SPSS. Immunoblotting results and response ratios in the devaluation experiment were analyzed using a 2-tailed, unpaired t-test. ANOVAs were used to compare response rates in other behavioral experiments, with repeating measures (RM) when appropriate. Tukey’s post-hoc tests were used in the case of significant interactions or main effects with >2 groups and are indicated in the figures. Values >2 standard deviations above the mean were considered outliers and excluded. For electrophysiological experiments, hyperpolarization was analyzed using a 1-sample t-test using 0 as a baseline. Rheobases before and during CNO were compared by a 2-tailed, paired t-test. The numbers of spikes in response to current steps were plotted and compared using a 2-way RM ANOVA followed by Bonferroni post-hoc tests.

Proportions of each dendritic spine subtype were generated by dividing a given spine subtype number by the total spine number for that dendritic segment. For dendritic spine subtype proportion and morphometric analyses, each animal contributed one value, reflecting the average of its dendrites. For initial subtype and morphometric comparisons, a 1-way ANOVA was used. Any significant effects were explored using Tukey’s post-hoc analyses. In the case of unequal variances, as analyzed by a Brown-Forsythe test, a Welch’s ANOVA was used instead, with Dunnett T3 post-hocs. All knockdown comparisons were analyzed using 2-tailed, unpaired t-tests. Welch’s correction was used in the case of unequal variances. All experiments were conducted in at least 2 independent cohorts of mice. Sample sizes were determined based on power analyses and prior similar experiments in our laboratory.

## Acknowledgments

We thank Dr. A. J. Koleske for the phosphotyrosine kinase antibody, Ellen P. Woon and H. Arrowood for assistance with dendritic spine reconstruction, Meghan Wynne for assistance with pilot experiments, and Dan C. Li for his thoughtful contributions to the manuscript. The work in the S.L.G. lab is supported by NIH T32 GM008602, F30 MH117878, F31 MH109208, R01 MH117103, and P50 MH100023. The Yerkes National Primate Research Center is supported by the Office of Research Infrastructure Programs/OD P51 OD011132. Research reported in this publication was also supported in part by the Emory University Integrated Cellular Imaging Core and Children’s Healthcare of Atlanta.

## Supplementary Information for

**Fig. S1.**
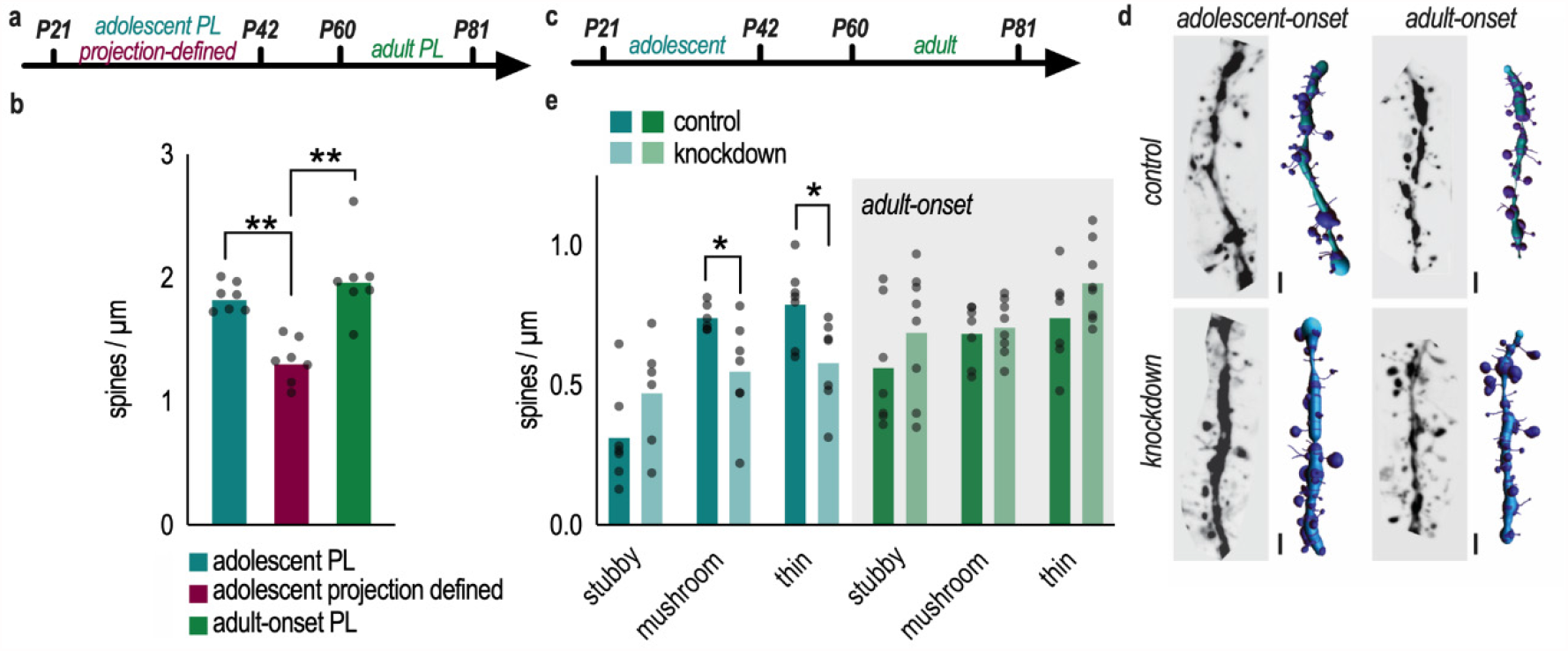
β1-integrin loss in adolescence, but not adulthood, decreases dendritic spine density. **a**. Timeline of experiments portraying time of viral vector delivery and euthanasia. **b**. Projection-defined PL neurons had lower dendritic spine densities than either mature or adolescent PL neurons that were not defined by projection [*F*_(2,18)_=17.10, *p*<0.0001]. **c**. Timeline of β1-integrin reduction. **d**. Adolescent-onset β1-integrin reduction decreased the density of mushroom- [*t*_(11)_=2.469, *p*=0.0312] and thin-type [*t*_(12)_=2.614, *p*=0.0210] dendritic spines. Adult-onset β1- integrin reduction did not affect dendritic spine densities. **e**. Representative dendrites, with their associated 3D reconstructions. Scale bar=2 μm. Bars=means, symbols=individual mice. \**p*<0.05, \*\**p*<0.001.

**Fig. S2.**
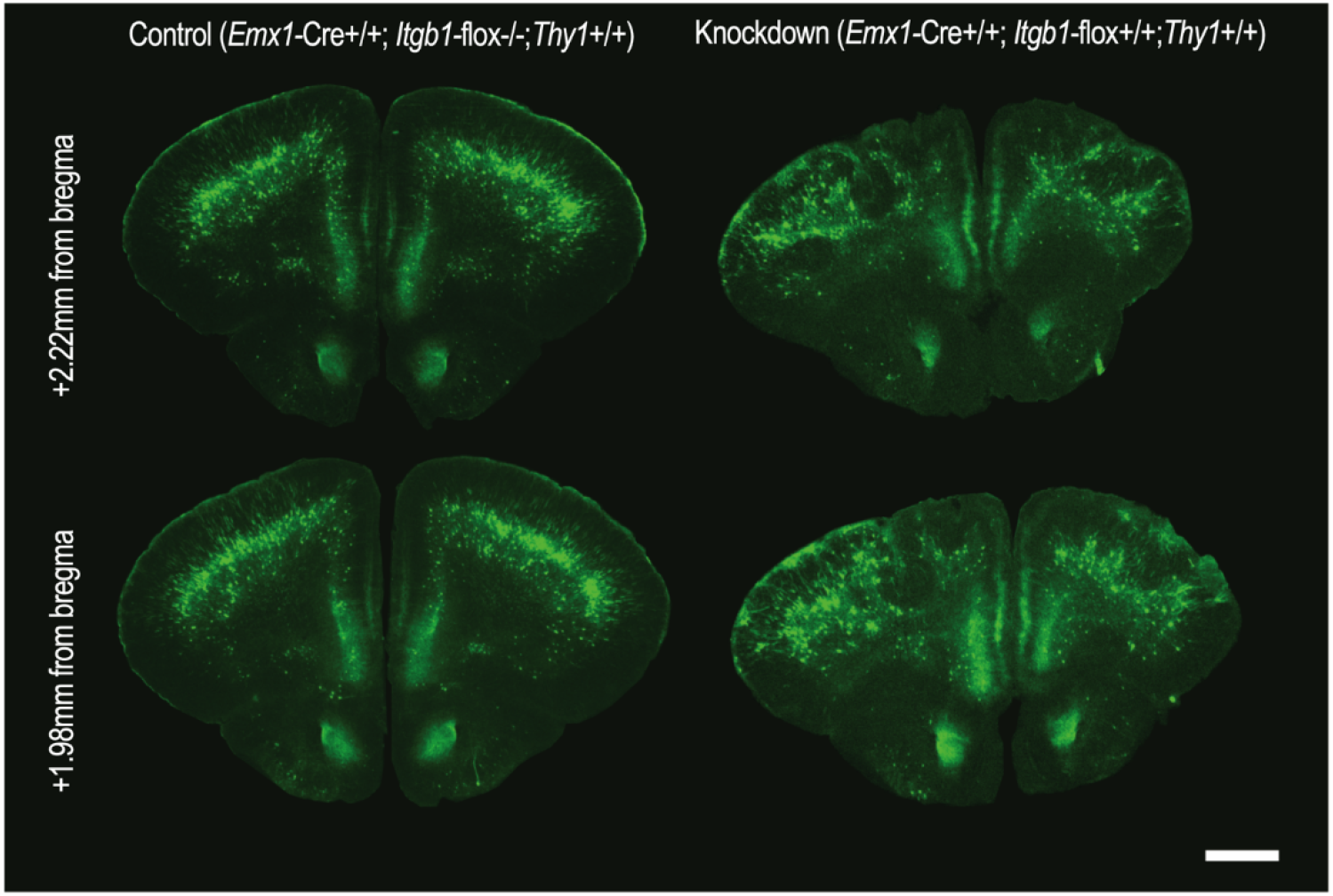
β1-integrins are required during embryogenesis for proper cortical layering. (left) Control brain sections at P21 display typical cortical organization, with layer V neurons expressing YFP. (right) Reduction of *Itgb1* starting at embryonic day 11.5 results in disorganized cortical layers at P21. Scale bar=1 mm.

